## Supplement for "Dynamic encoding of social threat and spatial context in the hypothalamus"

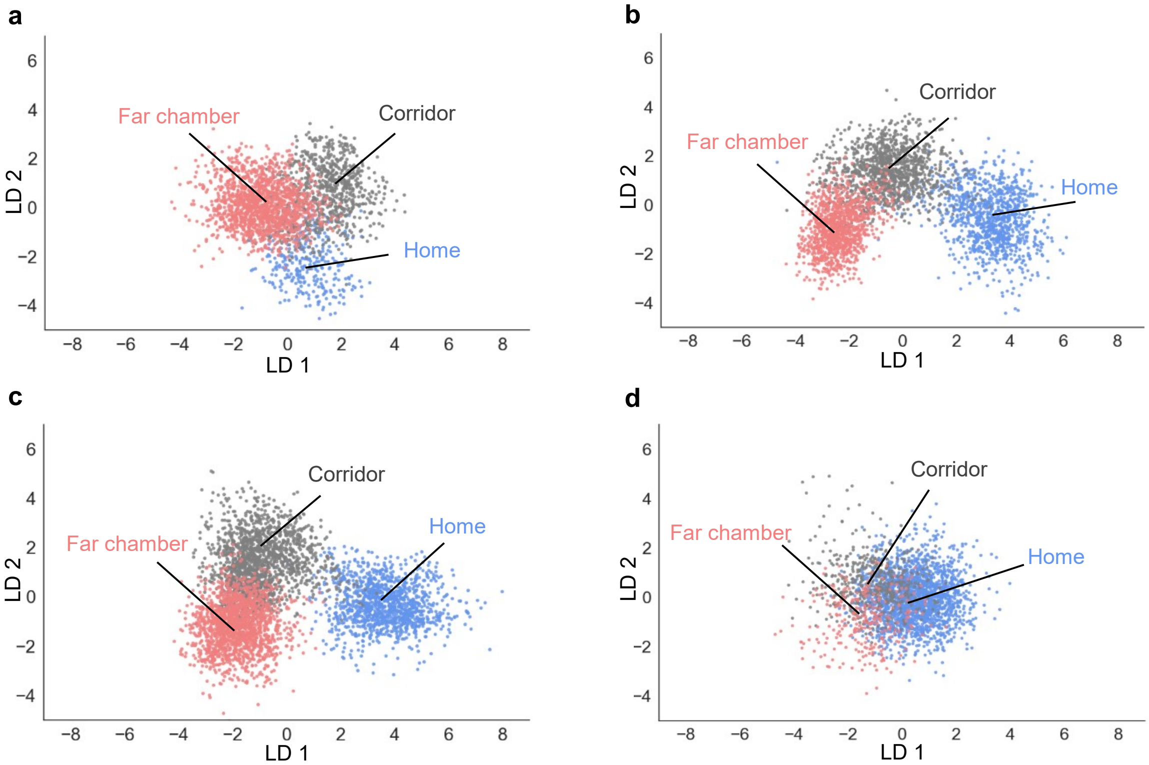


Krzywkowski et al., **Figure S1**

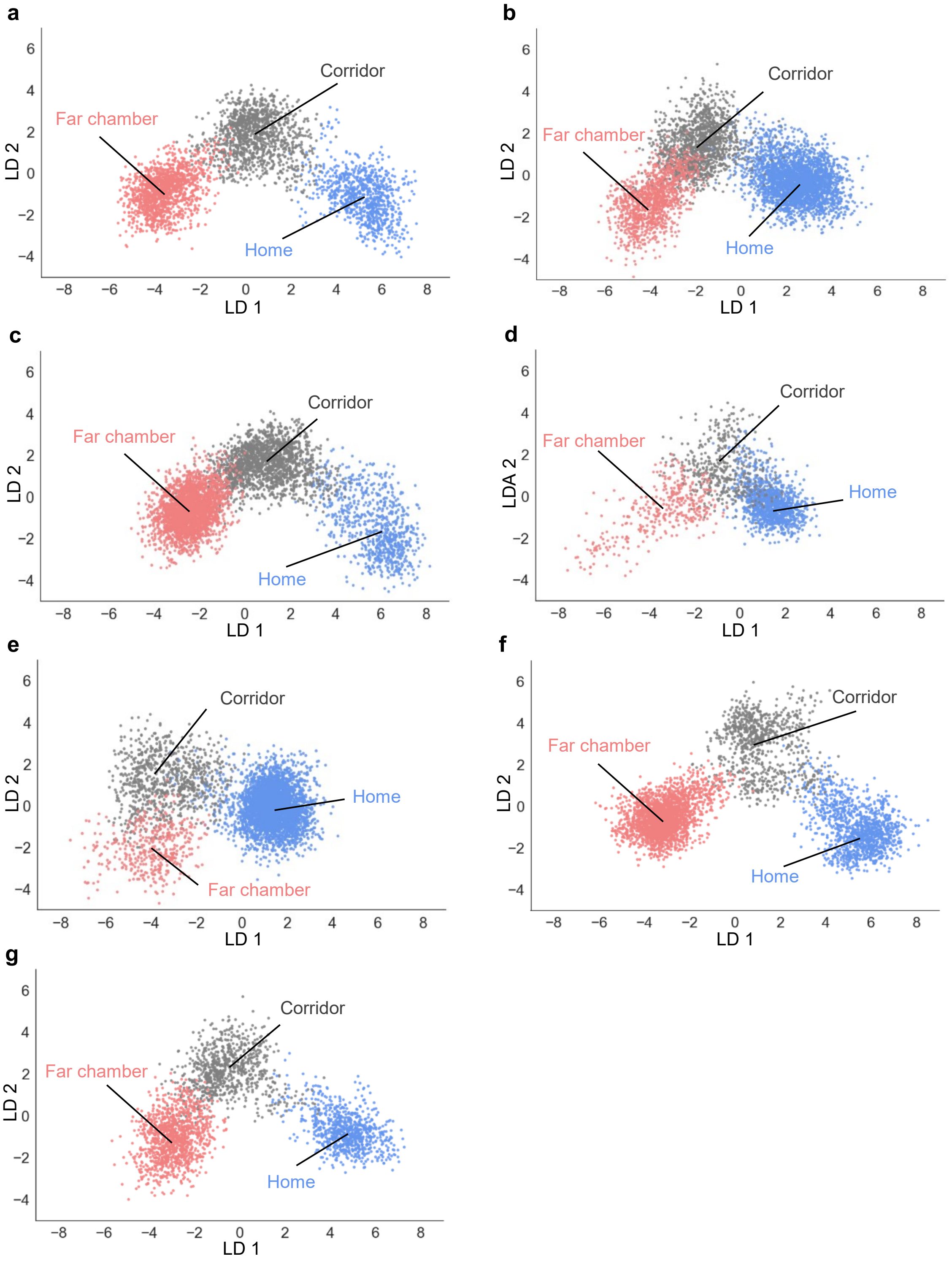


Krzywkowski et al., **Figure S2**

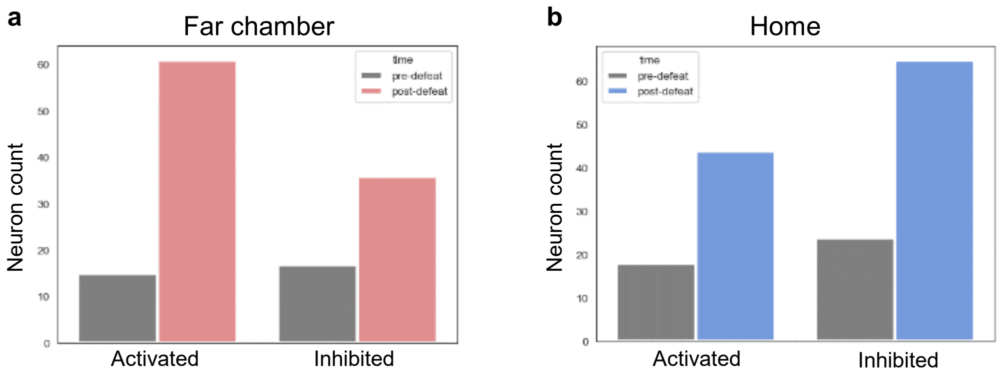


Krzywkowski et al., **Figure S3**

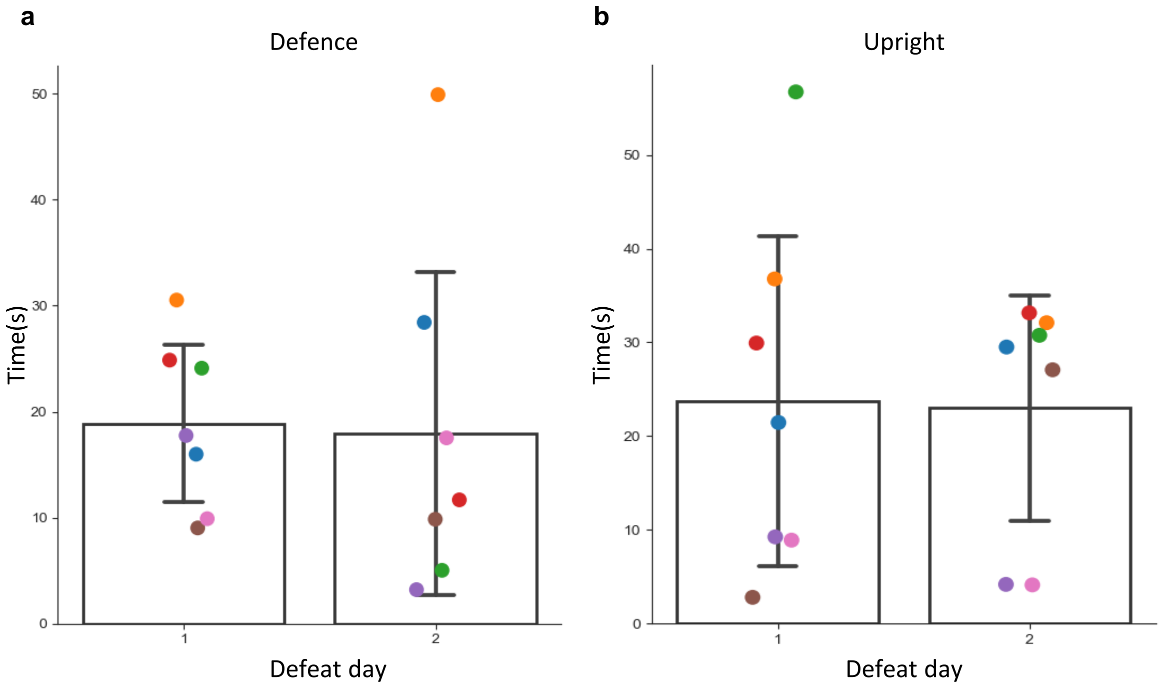


Krzywkowski et al., **Figure S4**

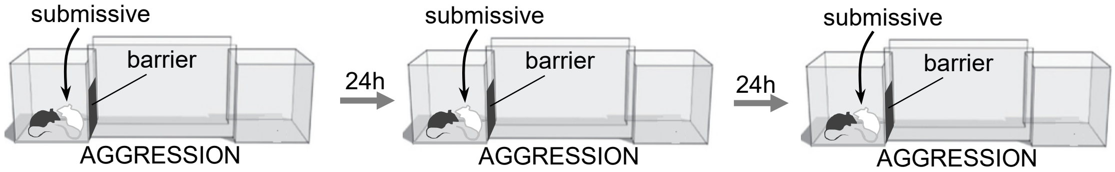


Krzywkowski et al., **Figure S5**

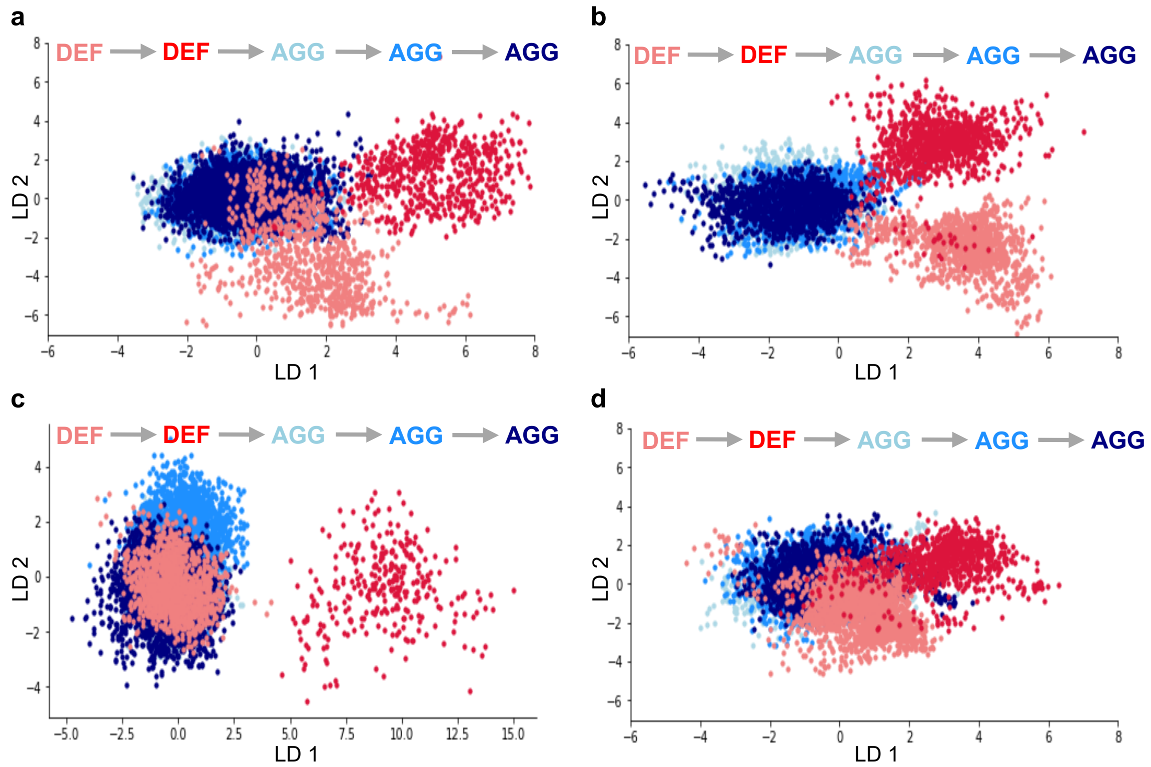


Krzywkowski et al., **Figure S6**

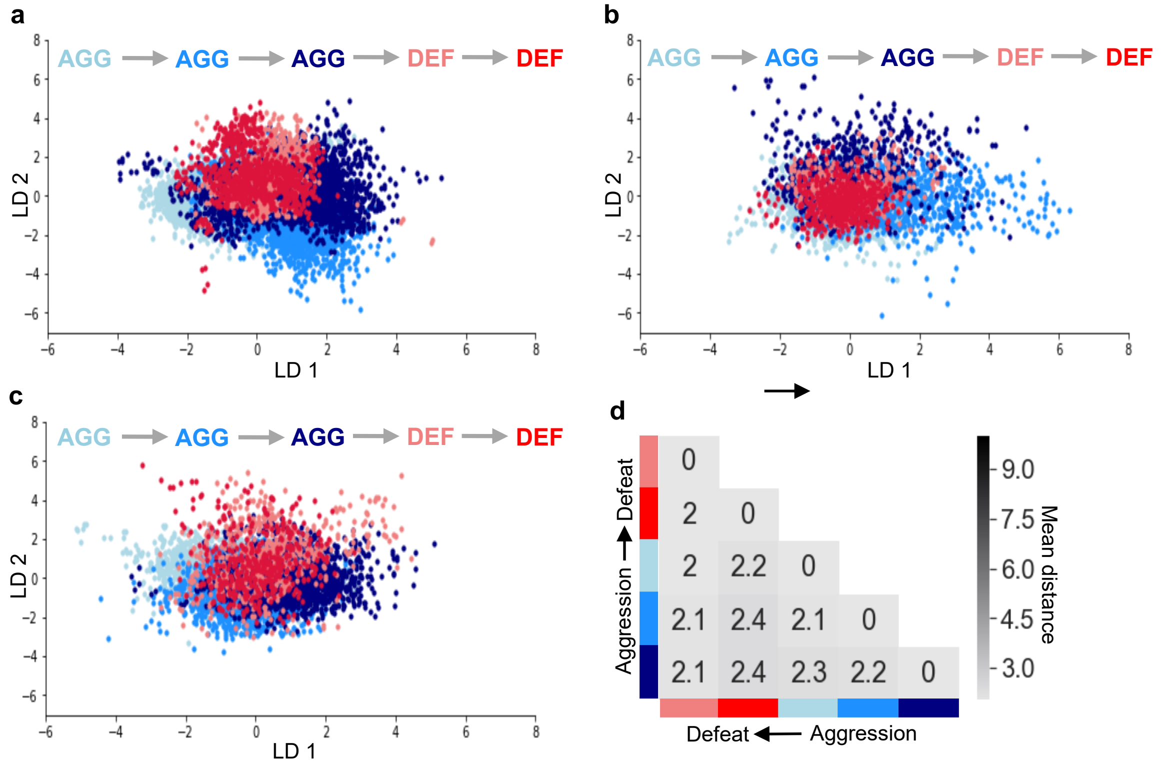
a
AGG → AGG → AGG → DEF → DEF
LD 2
LD 1
b
AGG → AGG → AGG → DEF → DEF
LD 2
LD 1
c
AGG → AGG → AGG → DEF → DEF
LD 2
LD 1
d
Aggression → Defeat
Defeat ← Aggression
Mean distance
0
2 0
2 2.2 0
2.1 2.4 2.1 0
2.1 2.4 2.3 2.2 0
9.0
7.5
6.0
4.5
3.0

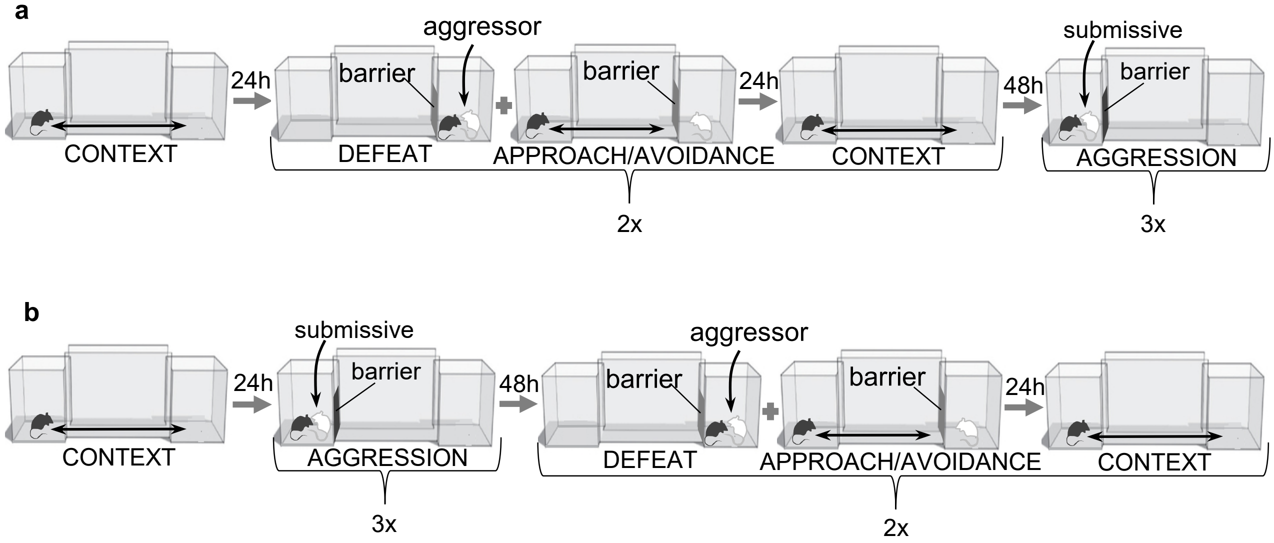


Krzywkowski et al., **Figure S8**

| Mouse | Assessment | Defense | Flight | Sniff | Attack | Territory |
|---|---|---|---|---|---|---|
| 1 | 41 | 39 | - | 55 | 55 | 44 |
| 2 | 57 | 58 | 58 | 61 | 61 | 61 |
| 3 | 43 | 30 | 44 | 46 | - | 44 |
| 4 | 59 | 60 | - | 71 | 72 | 62 |
| 5 | 35 | 36 | - | 41 | 41 | 35 |
| 6 | 53 | 53 | 53 | - | - | 54 |
| 7 | 38 | 43 | 43 | 36 | 37 | 43 |
| **Total** | **326** | **319** | **198** | **310** | **266** | **343** |

### Supplementary Figure Legends

***Figure S1.*** **Representation of territory in LD space *before* defeat for all mice.** (**a-d**) LDA plots of neuron ensemble activity for all mice in the home, far chamber, or corridor before defeat. Note that data for the mouse in panel **a** is the same as in **Figure 2m**. Each data point represents a frame of calcium imaging data projected onto the first two linear discriminants.

***Figure S2.*** **Representation of territory in LD space *after* defeat for all mice.** (**a-g**) LDA plots of neuron ensemble activity for all mice in the home, far chamber, or corridor after defeat. Note that data for the mouse in panel **a** is the same as in **Figure 2p** and panels **a-d** correspond to mice in **Figure S1a-d**, respectively. Each data point represents a frame of calcium imaging data projected onto first two linear discriminants.

***Figure S3.*** **Number of territory-responsive neurons increases after defeat.** Bar plot showing the number of positive (Activated) and negative (Inhibited) responding (0.35 > auROC > 0.65) neurons in the (**a**) far chamber and (**b**) home chamber before and after defeat (N = 4).

***Figure S4.*** **No difference between defensive behaviors on the two defense days.** (**a**) Plot showing time spend defending for all mice during defense days 1 and 2 (P > 0.05). (**b**) Plot showing time spend in upright position for all mice during defense days 1 and 2 (P > 0.05).

***Figure S5.*** **Aggression behavioral paradigm.** On the aggression day a BALB/c intruder was introduced into closed home chamber for 10 min after which the intruder was removed. Procedure was repeated for three days.

***Figure S6.*** **Representation of *forward* order defeat and aggression states in LD space for all mice.** (**a-d**) LDA plots of neuron ensemble activity for all mice during repeated defeat and aggression episodes. Inset indicates order of defeat and aggression episodes. Each data point represents a frame of calcium imaging data projected onto first two linear discriminants. Note that data for the mouse in panel **a** is the same as in **Figure 4a**.

***Figure S7.*** **Representation of *reverse* order defeat and aggression states in LD space for all mice.** (**a-d**) LDA plots of neuron ensemble activity for all mice during repeated defeat and aggression episodes. Inset indicates order of defeat and aggression episodes. Each data point represents a frame of calcium imaging data projected onto first two linear discriminants. Average distances between clusters of neuron ensemble data between defeat and aggression episodes for all mice during reverse order testing (N = 3; colors refer to episodes in **a-c**).

***Figure S8.*** **Forward and reverse behavioral paradigms.** Graphic depicting complete behavioral paradigm in (**a**) forward and (**b**) reverse order.

***Table S1.*** **Number of ROIs for each mouse that were assessed for the indicated behavior.** Territory includes both Home and Far chamber cells.
